# Direction-Specific Effects of Artificial Skin-Stretch on Stiffness Perception and Grip Force Control

**DOI:** 10.1101/2023.06.30.547233

**Authors:** Mor Farajian, Raz Leib, Hanna Kossowsky, Ilana Nisky

## Abstract

When interacting with an object, we use kinesthetic and tactile information to create our perception of the object’s properties and to prevent its slippage using grip force control. We previously showed that applying artificial skin-stretch together with, and in the same direction as, kinesthetic force increases the perceived stiffness. Here, we investigated the effect of the direction of the artificial stretch on stiffness perception and grip force control. We presented participants with kinesthetic force together with negative or positive artificial stretch, in the opposite or the same direction of the natural stretch due to the kinesthetic force, respectively. Our results showed that artificial skin-stretch in both directions augmented the perceived stiffness; however, the augmentation caused by the negative stretch was consistently lower than that caused by the positive stretch. Additionally, we proposed a computational model that predicts the perceptual effects based on the preferred directions of the stimulated mechanoreceptors. When examining the grip force, we found that participants applied higher grip forces during the interactions with positive skin-stretch in comparison to the negative skin-stretch, which is consistent with the perceptual results. These results may be useful in tactile technologies for wearable haptic devices, teleoperation, and robot-assisted surgery.

## I. Introduction

When interacting with objects, we receive feedback from multiple sensory modalities and integrate them to estimate the objects’ dynamics. From these estimations, we form our perception, e.g. our perception of stiffness, and create internal models that allow us to predict the consequences of the interactions [1].The stiffness of elastic objects is the linear relationship between the penetration into the object and the resulting force [2], [3].

Since we do not possess stiffness sensors, stiffness perception is formed at higher levels of the sensorimotor system by integrating the deformation of the object and the sensed force [4]. There are two major types of force sensing modalities in the human body: kinesthetic, which is sensed by muscle spindles and Golgi tendon organs, and tactile, which is sensed by cutaneous mechanoreceptors in the skin. The integration of the two modalities is important for estimating the mechanical properties of objects (e.g. stiffness, viscosity, and mass), and for generating internal representations of the object dynamics toward planning the necessary grip forces to apply [5], [6].

During tool-mediated interactions with objects, we apply grip force perpendicularly to the tool to prevent it from slipping due to different forces acting on it. It is well established that the grip force is coupled to the load force during interactions with elastic objects [7]. Grip force control involves two different mechanisms: predictive and reactive, both of which are vital for preventing the slippage of manipulated objects. The predictive grip force control is comprised of two components: (1) a baseline, which is applied when there is no load force acting and (2) a modulation of the grip force in anticipation of the load force [8]. The baseline provides a safety margin against slippage and depends on one’s certainty in the estimation of the object dynamics [9]. The modulation is adjusted in anticipation of the load force, and depends on an internal representation of the object dynamics [1], [10] and on the slipperiness of the finger interface [8], [11]. This modulation starts from the moment our fingers touch the object, and is updated during repeated interactions [12]. Several studies [13], [14] have used the grip force modulation in anticipation of the load force as an index of prediction in the control of movements. These studies revealed that the intended peak grip force can be predicted from the grip force and the rate of its change at the time of initial contact with the object [13], [14].

Understanding the processing of kinesthetic and tactile information in stiffness perception and grip force control can contribute to several fields, including the development of teleoperated technologies that display force information [15]– [17]. Due to stability issues, most of these technologies suffer from the absence of haptic information, or present haptic information that is of low gain and quality [18]. To overcome this challenge, studies have demonstrated other techniques to present force information in ways that do not affect the stability of the teleoperation system [19]–[22]. One technique that can be used to communicate the force information without affecting the stability of the teleoperation is through tactile feedback. Quek et al., [23] designed a device that provided tangential and normal artificial skin deformation for teleoperated surgical tasks. Schorr et al., [19] developed a wearable fingertip haptic device with the ability to render both shear and normal artificial skin deformation to the fingerpad. Abiri et al., [20] created a bimodal vibrotactile system that can be used with the da Vinci surgical robot.

Artificial skin-stretch has also been used to convey direction [24], augment friction [25], and replace kinesthetic information in navigation tasks [26]. While these, and other examples, demonstrate the benefits of artificial tactile feedback, the application of this feedback is still performed using empiric tuning and pilot studies. Furthermore, models of how artificial tactile information is processed by users are still limited.

Recent studies showed that applying artificial skin-stretch in the same direction as kinesthetic load force augments the perceived stiffness linearly to the amount of stretch [27]–[29]. Additionally, this skin stretch increases the predictive grip force modulation in anticipation of the load force [27]. Importantly, artificial tactile stimulation is added to the natural tactile stimulation that is applied during interactions with objects and kinesthetic haptic devices. In aperture-based skin-stretch devices [27], [28], this natural skin deformation is created as a result of the the contact between the fingers and the aperture when external forces are applied on the device. In all the studies referenced above, the artificial skin-stretch was applied in the same direction as the natural skin deformation caused by the kinesthetic forces. However, artificial skin-stretch applied in different directions may differ in its effect on perception and grip force control due to the different stimulation of the mechanoreceptors, and to their responses to these different stimulations.

The only study that examined the effect of artificial skin-stretch applied in the opposite direction to the kinesthetic force observed a large variability in the perceptual effects between participants [30]. Furthermore, the influence of negative artificial skin-stretch on grip force control is unclear. Hence, to reach a clear conclusion regarding the perceptual and grip force effects of artificial stretch stimulation, we now study the effect of positive (consistent with the natural) and negative (opposite to the natural) artificial skin-stretch. Moreover, we wish to support our experimental investigation with a mechanistic understanding of the effects by proposing a computational model that relies on the neural responses to the stimulation of our mechanoreceptors.

The mechanoreceptors in our skin play a crucial role in manipulation tasks – they directly provide information about mechanical interactions. Mechanoreceptor types differ from one another in the structure and the size of their receptive fields, in the densities within the separate sub-regions of the skin area, and in their adaptation rates [31]. Slowly adapting mechanoreceptors are more sensitive to steady skin deformation, whereas the rapidly adapting mechanoreceptors are more sensitive to transient motion on the skin, and alert the brain in case of slippage [31], [32]. The slowly adapting type 2 mechanoreceptors are the main mechanoreceptors that detect skin-stretch [33]. When interactions with objects lead to skin stretch deformation, these mechanoreceptors play a role in creating our perception of object motion direction and force.

Important to the current study, these mechanoreceptors exhibit a directional sensitivity; their discharge increases following stretch in one direction and decreases following stretch in other directions, providing highly discriminative information about the direction of skin-stretch [33]–[35]. The tactile information is coded by the firing rates of the afferent neurons, i.e., neurons that relay information from the mechanoreceptors to the nervous system. Tactile afferent neurons have higher firing rates due to stimulation in the preferred direction, and lower firing rates due to stimulation in the other directions [34], [36]. A weighted average of the preferred directions of a population of afferent neurons is called a population vector, which enables the estimation of force direction [34], [37].

In the current study, we use a similar experimental setup and methodologies to two of our previous works [27], [29]. Despite the technical similarities, the research questions addressed in these three studies are different. In our first work [27], we studied the effect of different magnitudes of positive artificial skin-stretch on stiffness perception and grip force control. We reported the magnitude of the increase in perceived stiffness and the timeline of its creation, as well as the effect on the predictive and reactive grip force control. Next, in [29], we studied if the perceptual augmentation caused by the positive artificial skin-stretch is impaired when visual displacement information is presented to the participants, and found that the presence of visual information weakens the augmentation. Hence, in both of these studies the artificial stretch was applied in the same direction as the natural stretch caused by kinesthetic forces.

Here, we conducted a stiffness discrimination experiments in which participants interacted with virtual objects, comprised of force feedback and artificial tactile skin-stretch applied in different directions. To further understand how artificial tactile stimulation affects perception and grip force control, we reproduce the results from [27] and extend this study to reveal the differences between the effects of positive and negative artificial skin-stretch on stiffness perception and grip force control. We used our setup and methods established in [27] to test our new research questions. We first focused on how the direction of the applied artificial skin-stretch affected stiffness perception. As the negative artificial stretch is applied in the opposite direction to natural skin-stretch, we hypothesized that it would cause a decrease in the perceived stiffness (that is, cause participants to underestimate the perceived stiffness). However, if participants would interpret the artificial tactile stimuli as an indication of a more slippery contact surface at the interface with the fingers, and not as a stretch, we would not expect to find a difference in the perceived stiffness between the two different directions of artificial stretch. In addition, we hypothesized that the negative skin-stretch may also increase the perceptual uncertainty experienced by the participants.

Thereafter, we examined the effect of positive and negative artificial skin-stretch on grip force control. On the one hand, if the perception and the control of grip force share similar stiffness estimation mechanisms, and if the negative artificial stretch would decrease the perceived stiffness, we would expect participants to apply a lower grip force modulation during trials with negative skin-stretch compared to trials with positive skin-stretch. On the other hand, if the negative artificial skin-stretch would increase the uncertainty experienced by the participants, we would expect to find an increase in the grip-force baseline during trials with negative skin-stretch.

A preliminary version of this study was reported in [38], in which we performed a pilot study with N=4 participants. In the current study, we modified the experimental protocol, increased the sample size to achieve a sufficiently powered study, and proposed a computational model based on a neural population vector with a bimodal distribution of preferred directions that explains the perceptual results.

## II. Methods

### A. Participants

40 right-handed participants (19 females and 21 males, aged between 21-27, average age: 24.58 ± 1.40) completed the experiment. Participants signed a written informed consent form prior to the experiment after hearing an explanation by the experimenter. The procedures and the consent form were approved by the Human Subjects Research Committee of Ben-Gurion University of the Negev, Be’er-Sheva, Israel, approval number 1283-1, dated July 6th, 2015. The participants were compensated for their participation, regardless of their success or completion of the experiment.

### B. Experimental Setup

The experimental setup used in this work is identical to the one we designed in [27]. Participants interacted with a virtual environment using a skin-stretch device that was mounted on a PHANTOM® Premium 1.5 haptic device (3D SYSTEMS). The haptic device generated load force feedback and natural skin-stretch, and the skin-stretch device generated the additional artificial skin-stretch stimulation. The direction and the magnitude of the load force and the artificial skin-stretch in each trial were controlled using the Open Haptics API, written in C++ (Visual Studio 2010, Microsoft). The participants viewed the virtual environment through a semi-silvered mirror that blocked their view of their hand and showed the projection of an LCD screen above it (Fig. 1(b)), and wore noise cancelling headphones (Bose QC35) to eliminate auditory cues from the motor of the skin-stretch device.

**Fig. 1.**
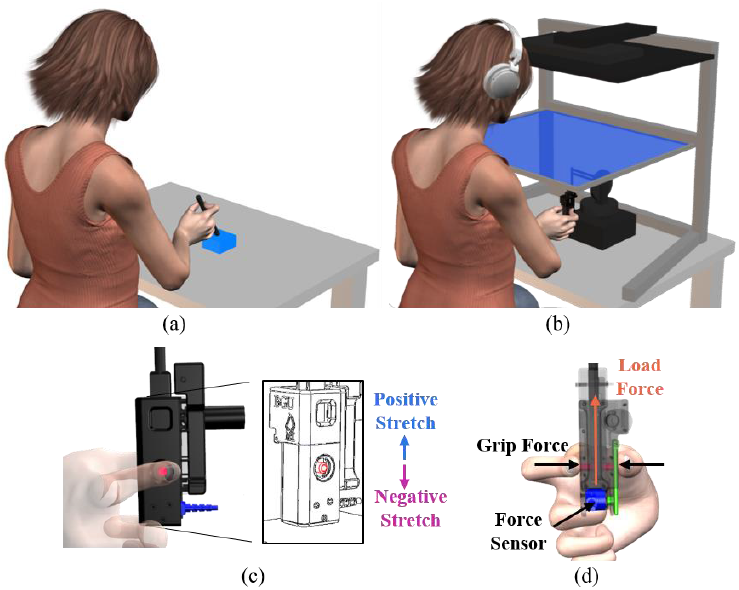
Experimental system. (a) Our experimental setup was designed to emulate tool-mediated interactions with real objects, akin to touching a sponge with a stick. (b) The participants sat in front of a virtual reality rig, and held the skin-stretch device, which was mounted on the end of a haptic device, both rendering interaction with a virtual object. (c) Side view of the skin-stretch device. The participants used the thumb and index finger of their right hand to grasp the device and folded their other three fingers. Two tactors (red rod) came into contact with the skin of the fingers and moved in the vertical direction to stretch the skin through tactor displacement. The positive artificial skin-stretch was applied in the upward direction (blue arrow), and the negative artificial skin-stretch was applied in the downward direction (purple arrow). (d) Back view of the skin-stretch device. The load force was applied by the haptic device in the upward direction and the grip force is the perpendicular force between the digits and the object. A force sensor (blue) was embedded in the device to measure the grip force that the participants applied, via the lever (green), which transmitted the grip force from the contact point to the sensor.

In our skin-stretch device, two skin-stretch tactors (rubber Lenovo Trackpoint Classic dome with a rounded surface and a rough sandpaper-like texture) were attached to a vertical bar. Around the tactors, on the outer shell of the device, there were two round apertures on which participants placed their thumb and index fingers to grasp the device. Participants were asked to place their fingers horizontally (parallel to the table). This setup caused the fingerpads to press on the tactors, and when the vertical bar moved, it caused the skin of the fingerpads holding the device to be artificially stretched in the vertical direction by the tactors. The positive artificial skin-stretch was applied in the upward direction (the same direction as the load force and the natural skin-stretch), and the negative artificial skin-stretch was applied in the downward direction (the opposite direction to the load force and the natural skin-stretch) (Fig. 1(c) and 1(d)).

We measured the grip force applied by the participants using a force sensor (ATI, Nano17), which was mounted on the lower end of the device such that participants did not place their fingers directly above it (Fig. 1(d)). The left side of the outer shell consisted of a ‘door’ with an axis on its upper end, and a cylindrical protrusion facing the force sensor. When the device was held with the index finger and thumb on the apertures, the protrusion pressed the force sensor, and the relative grip force was measured. The skin-stretch device with the embedded force sensor was identical to the one described in [27]. Because of the colocation of the tactor movement mechanism and the ideal placement of the force sensor, we could not measure the grip force directly. However, the division of the grip force between the tactor and the aperture, and the placement of the force sensor at a distant location, allowed (through the law of conservation of angular momentum) measurement of a downscaled version of the applied grip force. That is, we measured trends in the grip force that were proportional to the actual grip force that the participants applied.

The load force and skin-stretch were proportional to the vertical position of the end-point of the haptic device and were applied only when participants were in contact with the virtual object, which was defined to be the negative half of the vertical axis of the robot’s coordinate frame:

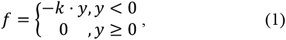

where k [N/m] is the stiffness, and y [m] denotes the participant’s hand position.

We used the skin-stretch device to apply tactile stimuli by means of tactor displacement:

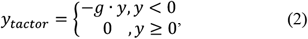

where *g* [mm/m] is the tactor displacement gain and y [m] is the hand position. The artificial skin-stretch was applied either in the same direction as (*g* = 80 *mm*/*m*), or in the opposite direction to (*g* = −80 *mm*/*m*) the applied kinesthetic load force.

### C. Protocol

In a forced-choice stiffness discrimination task, participants made downward vertical probing movements into pairs of virtual objects that applied kinesthetic and tactile feedback, and reported which of the two had a higher level of stiffness. The virtual elastic objects were designated *standard* and *comparison*, and were indicated to the participants by the color of the LCD screen, which was pre-defined pseudo-randomly to be either red or blue (Fig. 1(b)). That is, each object could be either red or blue in each trial, and the order of the presented colors was defined pseudo-randomly. In each trial, participants probed the first virtual object four times, and then raised the end-point of the haptic device to at least 3 cm above the boundary of the virtual object to switch to the second object. They then probed the second virtual object four times, and reported which object felt stiffer by pressing a keyboard key with the color corresponding to the screen color of the stiffer object. To begin the next trial, participants again raised the end-point of the haptic device to at least 3 cm above the boundary of the virtual object.

The experiment was divided into two sessions that were completed over two days. The participants were randomly split into two groups. Group 1 (N=20) completed the positive stretch session on the first day and the negative stretch session on the second day, and Group 2 (N=20) completed the two sessions in the opposite order. In each session, there were two experimental conditions: (1) only force feedback (control) and (2) force feedback with positive/negative artificial skin-stretch.

Throughout the experiment, the *comparison* virtual object applied only load force feedback, and the *standard* virtual object applied load force, and in some trials, also applied artificial tactile skin-stretch. The *standard* virtual object always had a stiffness value of 85 N/m, and in trials with artificial skin-stretch stimulation the skin-stretch gain was +80 mm/m in the positive stretch session, and -80 mm/m in the negative stretch session. This artificial skin-stretch gain was chosen because in our previous studies [27], [29] a clear augmentation in the perceived stiffness was demonstrated due to this level of positive artificial stretch. The stiffness level of the *comparison* virtual object was selected in each trial from a range of 12 values, evenly spaced between 30-140 N/m.

The participant began each session with 24 training trials to become familiarized with the experimental setup. During the training, both virtual objects applied only load force feedback, and at the end of each trial participants received feedback of ‘right’ or ‘wrong’ on their response. The remaining trials in each session were the test trials. They each contained 12 *comparison* stiffness levels and two *standard* conditions, amounting to a total of 24 *standard*-*comparison* pairs, each of which was repeated eight times throughout the experiment, resulting in 192 test trials. The order of the trials within each session was pseudo-randomized prior to the experiment. The duration of each session was about 40 minutes.

### D. Data Analysis

#### 1. Stiffness Perception

For each of the 40 participants, we used the Psignifit toolbox 2.5.6 (see http://bootstrap-software.org/psignifit/) [39] to fit psychometric curves to the probability of responding that the *comparison* virtual object was stiffer than the *standard* as a function of the difference between the stiffness levels of the two virtual objects. We repeated this procedure for the four experimental conditions (two force only conditions, one positive stretch condition and one negative stretch condition), and computed the point of subjective equality (PSE) and the just noticeable difference (JND) of each psychometric curve. The PSE is the stiffness level at which the probability of responding comparison is half. In this work, we chose to analyse the ΔPSE, which is the difference between the PSE and the stiffness value of the *standard* virtual object (85 N/m). Hence, a positive ΔPSE value indicates a rightward shift of the psychometric curve and an overestimation of the *standard* virtual object stiffness, whereas a negative ΔPSE value indicates an underestimation of the *standard* virtual object stiffness. The JND quantifies the sensitivity of the participants to small differences between the stiffness levels of the two virtual objects, and is an indication of the uncertainty experienced by the participants when choosing which virtual object was stiffer.

#### 2. Grip Force Control

We recorded the grip force data and filtered it using the MATLAB function *filtfilt* with a 2^nd^ order Butterworth low-pass filter, with a cutoff frequency of 12Hz, resulting in a 4^th^ order filter, with a cutoff frequency of 9.62Hz. We examined the grip force applied by the participants for every trial in each of the four different stretch conditions.

During trials with artificial skin-stretch, the grip force measurement is distorted because of the movement of the tactors (see [27] for more information). Hence, it was not possible to use the actual maximum grip force applied during these trials. To analyze the maximum grip force during trials with artificial skin-stretch, we followed the approach proposed in [13], [14], where grip force signals were mechanically disturbed by the impact loads following collisions. It is well documented that an increase in the grip force, and the rate of its change with respect to time, precedes the increase in load force during contact with dynamic objects [40], [41]. Therefore, a multivariate linear regression model can be used to predict the intended peak grip force from the value of the grip force and its rate at the point of initial contact with the object [13], [14]. We previously validated this approach with our setup in [27].

In [27] we used stretch-catch probes (that is, probes in which we maintained the load force but omitted the skin-stretch) to train a multiple regression model that will predict the peak grip force from the value of the grip force and its rate at the point of initial contact with the object. As this experiment did not contain stretch-catch probes, we used the model we presented in [27]. The experiments were performed using the same system, and the experimental protocols of the two studies were similar. In addition, the coefficients we received in [27] were similar to those described in [13], [14]. Therefore, we believe it is valid to use the model we found in our previous study [27] to predict the intended grip force peak in this study. The *grip force at contact* was calculated as the grip force at the first sample in contact with the elastic virtual object. The grip force rate at contact was calculated with a backward difference approximation of the derivative of the grip force signal with respect to time. *The grip force rate at contact* represents the grip force trend during the entrance into the virtual object. In some of the probing movements we observed negative grip force rate at contact, indicating a decrease in the grip force or a large phase shift between the load and the grip force. It is well documented that the grip force and load force signals are coupled together during interactions with elastic virtual objects [7]. Therefore, we expect that an increase in the load force signal will lead to an increase in the grip force signal. We do not have a conclusive explanation as to why the reaction to skin-stretch would be an initial decrease of the applied grip force. One possibility is that the decrease was the result of an unpleasant or painful interaction, however, none of our participants indicated any discomfort. Because the study was designed predominantly around perception, we could not further investigate the origin of these negative grip force rates. Instead, we used a conservative approach, and excluded from the analysis probing movements in which the rate was negative. As a result, six participants were completely excluded from the grip force analyses because in all of their trials the rate was negative. The resulting model was:

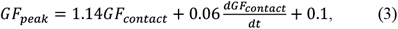

with *R*^2^=0.732, and both independent variables contributing significantly to the prediction of the *predicted grip force peak* (p<0.001).

We used this model to quantify the evolution of the *grip force modulation* in anticipation of the load force. The *predicted grip force peak* is determined by the *grip force at contact*, as well as by the modulation of grip force in anticipation of the load force. To get a better assessment of the evolution of the *grip force modulation* in anticipation of the load force, we calculated the *grip force modulation* by subtracting the *grip force at contact* from the *predicted grip force peak*. In addition, we calculated the *grip force baseline* as the deepest value of grip force between consecutive movements. Finally, to isolate the effect of artificial stretch stimulation on each of these grip force metrics, we calculated the difference between the values obtained due to the positive and the negative artificial stretches and each of their controls.

### E. Statistical Analysis

We examined the effects of the four different conditions on the ΔPSE and JND values across all the participants using a repeated-measures General Linear Model with the MATLAB statistic toolbox. The dependent variables in the two separate analyses were the ΔPSE and JND values. The independent variables were the stretch condition (1. no stretch of the negative session (CN), 2. negative stretch (N), 3. no stretch of the positive session (CP), and 4. positive stretch (P), categorical, 4 levels, df=3), and the participants (random, df=39). To compare between the different conditions, we performed four planned t-tests using the Holm-Bonferroni correction for multiple comparisons (Control-N vs. Negative, Control-P vs. Positive, Control-N vs. Control-P, and Negative vs. Positive). We presented the p-values after this correction (*p*_*corrected*_), and therefore the threshold significance level following the correction was 0.05.

To test the significance of the the effect of artificial stretch stimulation on the Δ*grip force baseline*, the Δ*predicted grip force peak*, and the Δ*grip force modulation*, and between the four probing movements, we fit separately a repeated-measures General Linear Model to each of the dependent variables, using the MATLAB statistics toolbox. The dependent variables were the differences between the values obtained due to the positive and negative artificial stretches and each of their controls. The independent variables were the stretch condition (categorical, 2 levels: 1. negative stretch, 2. positive stretch, df=1), the probing movement (categorical, 4 levels, df = 3), and the participants (random, df=33). The model also included the interaction between the ‘stretch condition’ and the ‘probing movement’ independent variables.

Next, to compare between the probing movements, we performed planned t-tests using the Holm-Bonferroni correction for multiple comparisons. When the interaction between the ‘stretch condition’ and the ‘probing movement’ factors were statistically significant, we performed the planned t-tests separately for each of the stretch conditions (positive and negative). However, when only the ‘probing movement’ factor was statistically significant, the planned t-tests were performed on the two stretch conditions together. The three planned t-tests we chose to perform were: first vs. second probing movements, second vs. fourth probing movements, and first vs. fourth probing movements. We chose these three comparisons based on our previous work [27], in which we showed that following exposure to artificial skin-stretch, participants immediately (after the first interaction) increased their grip force baseline, and gradually (until roughly the fourth interaction) increased the predictive *grip force modulation* in anticipation of the load force. We presented the p-values after this correction (*p*_*corrected*_), and therefore the threshold significance level following the correction was 0.05.

To assess whether the order of the sessions (positive or negative first) influenced the effects of the stretch on perception and on the control of grip force, we used a nested General Linear Model, where the participants variable was nested in the group-number variable (categorical, df=1).

## III. Results

### A. Order of Sessions

First, we examined the effect of the order of the sessions on the results. We did not find any significant effects of order on perception (rm-General Linear Model, PSE: main effect of ‘group number’: *F*_(1,38)_ = 1.03, *p* = 0.3162; JND: main effect of ‘group number’: *F*_(1,38)_ = 0.82, *p* = 0.3702). Similarly, we did not find any significant effects on the control of grip force (rm-General Linear Model, *Grip force baseline*: main effect of ‘group number’: *F*_(1,32)_ = 0.15, *p* = 0.7013; *Predicted grip force peak*: main effect of ‘group number’: *F*_(1,32)_ = 0.63, *p* = 0.4324; *Grip force modulation*: main effect of ‘group number’: *F*_(1,32)_ = 1.00, *p* = 0.3240). Therefore, we combined the two order groups for the remaining analyses in this paper.

### B. Stiffness Perception

At the population level, the addition of artificial skin-stretch, in the same or in the opposite direction as the kinesthetic load force, caused participants to overestimate the stiffness of the *standard* virtual object. However, we found that the perceived stiffness due to the negative stretch was consistently lower than that caused by the positive stretch.

The psychometric curves of a typical participant are presented in Fig. 2(a). The light blue and the light purple psychometric curves represent conditions without artificial skin-stretch, and the dark blue and dark purple curves represent conditions with artificial skin-stretch. In conditions without artificial skin-stretch, the PSE was close to zero. This means that the participant could accurately distinguish the stiffness levels of the two virtual objects. Stretching the skin of the typical participant in the opposite direction to that of the kinesthetic load force caused a rightward shift of the curve and a positive PSE, indicating that this participant overestimated the stiffness of the *standard* virtual object due to the artificial stretch. Stretching the skin of this participant in the same direction as that of the kinesthetic load force caused an even larger rightward shift of the PSE.

**Fig. 2.**
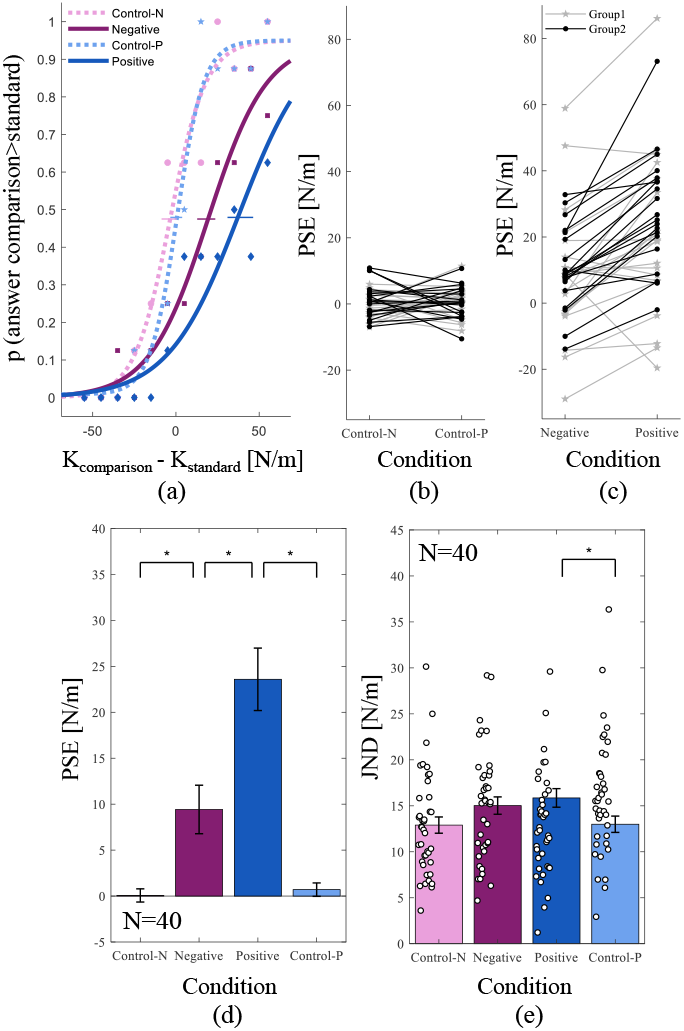
The effect of artificial skin-stretch in both directions on stiffness perception. (a) Example of psychometric curves of a typical participant for the different conditions (1. no stretch of the negative session, 2. negative stretch, 3. no stretch of the positive session, and 4. positive stretch). The abscissa is the difference between the stiffness levels of the comparison and the standard force fields, and the ordinate is the probability of responding that the comparison force field had a higher level of stiffness. The horizontal lines represent the standard errors for the ΔPSE values. (b) The ΔPSE values of all the participants in the two control conditions without the skin-stretch. (c) The ΔPSE values of all the participants in the two conditions with the artificial skin-stretch. The gray stars and lines represent the data of Group 1 (N=20, positive stretch session first), and the black circles and lines represent the data of Group 2 (N=20, negative stretch session first). (d) The averaged ΔPSE values across all the participants (N=40), as a function of the different conditions. (e) The JND values as a function of the different conditions. The black circles represent the data of each of the participants (N=40), and the colored bars show the average values across all the participants. The black error bars represent the standard errors of the estimated means, and the asterisks indicate a statistically significant difference (p<0.05).

Fig, 2(b) and 2(c) present the PSE values of each of the participants in the four different conditions. Both positive and negative artificial skin-stretche affected participants’ stiffness perception; our results show a wide range of underestimation and overestimation effects on the perceived stiffness [Fig. 2(c)]. For most of the participants (33 out of 40), both directions of artificial skin-stretch led to an increase in the perceived stiffness. Additionally, we found that the perceived stiffness in the positive stretch condition was generally higher than the perceived stiffness in the negative stretch condition; this was also true for the cases in which the artificial skin-stretch led to an underestimation in the perceived stiffness.

The colored bars in Fig. 2(d) show the PSE average results of all the participants. We found that both directions of artificial skin-stretch increased the perceived stiffness (PSE, rm-General Linear Model, main effect of ‘stretch condition’: *F*_(3,117)_ = 34.83; p < 0.0001). The planned t-tests confirmed that artificial skin-stretch augmented the perceived stiffness when it was applied in the same direction as the kinesthetic force (*t*_P−CP (117)_ = 8.71, *p*_corrected_ < 0.0001), consistently with [28], [27], as well as when it was applied in the opposite direction to the kinesthetic force (*t*_N−CN (117)_ = 3.56, *p*_corrected_ = 0.0011). In addition, the positive skin-stretch caused a greater augmentation effect relative to the augmentation effect caused by the negative skin-stretch (*t*_P−N (117)_ = 5.39, *p*_corrected_ < 0.0001).

The JND values of each of the participants, and their means, in the four different conditions, are depicted in Fig. 2(e). Even though the ‘stretch condition’ was statistically significant (JND, rm-General Linear Model, main effect of ‘stretch condition’: *F*_(3,117)_ = 5.55; p = 0.0013), the planned t-tests revealed that there was a significant difference only between the positive skin-stretch and its control (*t*_P−CP (117)_ = 3.22, *p*_corrected_ = 0.0065). That is, the positive skin-stretch led to an increase in the JND values relative to the JND in the condition without the skin-stretch.

### C. Bimodal Preferred Direction Distribution

To explain the perceptual results, we proposed a computational model that was based on a non-uniform distribution of the mechanoreceptors preferred direction. The model was based on the finding that mechanoreceptors have a preferred direction of skin-stretch stimulation [34], [36]. Each mechanoreceptor was represented by a vector in a specific direction, indicating the preferred direction of that mechanoreceptor. That is, when the direction of the skin-stretch coincides with the mechanoreceptor’s preferred direction, the firing rate of this mechanoreceptor is highest, while it is lowest in the opposite direction [34], [36], [42]. For simplicity, we do not distinguish between the directional sensitivity differences along the entire hierarchy of signal transduction from the receptors to the brain. Therefore, we model a simple case of a population of neural computation units with preferred direction distributions.

In the population level, the distribution of the preferred direction is non-uniform with more mechanoreceptors tuned to specific direction compared to other directions. To capture this characteristic, we suggested a bimodal distribution with two different and uneven peaks in two dominant directions: the first was the direction of the natural skin-stretch and the positive artificial skin-stretch, and the second was the direction of the negative artificial skin-stretch. The natural and positive artificial stretch were applied upwards in the same direction, while the negative artificial skin-stretch was applied in the downward direction, that is, at an angle of 180° relative to the natural and positive artificial stretch (Fig. 3(a)). We assumed that the positive direction (upwards) was more dominant than the negative direction (downwards), and implemented this dominancy by defining a smaller standard deviation for the positive direction distribution relative to that of the negative direction (Fig. 3(b)). To simulate the populations of neurons, we randomly drew vectors with preferred directions defined by the distributions of the positive and negative stretch distributions (Fig. 3(c)). Therefore, when drawing random vectors, the probability of drawing a vector with a preferred direction close to the direction of the positive stretch was greater than the probability of drawing a vector with a preferred direction close to that of the negative stretch.

**Fig. 3.**
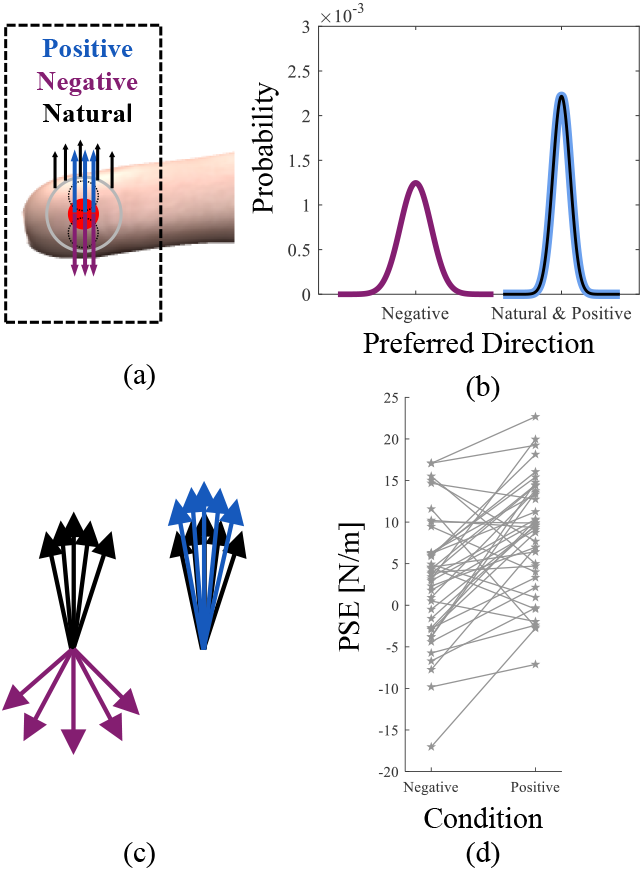
The proposed computational model for the effect of natural, positive artificial, and negative artificial skin-stretch stimulations of mechanoreceptors on stiffness perception. In the simulation of the model, we randomly selected the preferred directions of the mechanoreceptors to be affected in each trial. (a) The direction of the stretch stimulations during the experiment. The solid gray circle is the aperture on which participants placed their finger, and the red circle is the skin-stretch tactor. The dotted black circles are the range in which the skin-stretch tactor could move; the upper circle represents the positive stretch and the bottom circle represents the negative stretch. The blue upward arrows and the purple downward arrows represent the positive and negative artificial stretches, respectively, and the black upward arrows represent the natural stretch due to the load force applied by the robotic device. (b) The distribution of the preferred directions of a hypothetical population of mechanoreceptors is bimodal with uneven peaks in the two dominant directions: positive and negative. The black and blue distributions are the probabilities of affecting a mechanoreceptor with a preferred direction that is close to the direction of the natural and positive artificial stretch, respectively. The purple distribution is the probability of affecting a mechanoreceptor with a preferred direction that is close to the direction of the negative artificial stretch. (c) An example of the preferred directions of mechanoreceptors that were randomly chosen to be affected in the artificial negative (left) and artificial positive (right) conditions. (d) The results of the simulation: the calculated PSE values across all the simulated repetitions (N=40), as a function of the negative and positive artificial stretch conditions (compared to our experimental results in Fig. 2(C)).

Based on the bimodal preferred direction distribution, we simulated the two experimental stretch conditions we used in this study: positive and negative artificial skin-stretch. For each stretch condition we drew 10 mechanoreceptors with different preferred directions: five for the natural stretch and five for the artificial stretch. The five mechanoreceptors of the artificial stretch were stimulated by the skin-stretch tactor (the red rod in Fig. 3(a)), and the five mechanoreceptors of the natural stretch were stimulated by the aperture surrounding the tactor (represented by the solid grey circle in Fig. 3(a)). The natural skin-stretch mechanoreceptors were stimulated at the contact between the fingers and the aperture due to the application of the kinesthetic load force, regardless of the tactor movement (the black arrows in Fig. 3(a)). Therefore, the natural skin-stretch mechanoreceptors provided the same force information in the upward direction in both the positive and the negative artificial stretch conditions. On the other hand, the artificial skin-stretch mechanoreceptors were stimulated as a result of the movement of the skin-stretch tactors (the blue and purple arrows in Fig. 3(a)). The positive artificial stretch therefore provided force information in the upward direction, and the negative artificial stretch provided force information in the downward direction.

We simulated the two experimental stretch conditions according to the following steps:

For the natural stretch and the positive artificial stretch, we drew five mechanoreceptors for each stimulation according to equation 4, and for the negative artificial stretch we drew five mechanoreceptors according to equation 5 (Fig. 3(b)):

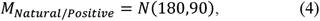

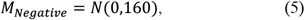

where *N* represents a normal distribution. To take into account the periodicity of the angle, we corrected the drawn mechanoreceptors with preferred direction to be within a range of 360° around the mean.

We simulated the responses of these random mechanoreceptors to stretch in the vertical direction. The magnitude of the responses of mechanoreceptors with preferred directions coinciding with those of the stretch stimulus was equal to 1. The magnitude of the responses of mechanoreceptors with preferred direction that did not coincide with the stretch stimulus was equal to the cosine of the angle between the preferred direction and the direction of the stretch:

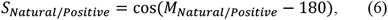

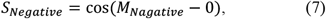

That is, a preferred direction that is closer to the direction of the stretch stimulus will lead to a response of greater magnitude.

We assumed a linear transformation between stretch and force information; that is, the stretch information, which was represented by the magnitude of the preferred direction, was proportional to the magnitude of the force for each mechanoreceptor.

After calculating the individual preferred directions, we calculated the population vector [43] as a weighted average of the individual preferred direction vectors:

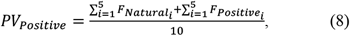

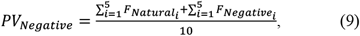

Where *F* [*N*] is the estimated force. This calculation provided us with a population vector whose magnitude was proportional to the general force information given by the entire population of mechanoreceptors. Finally, we calculate the PSE values in units of stiffness:

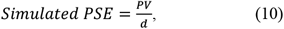

where *d* is the average penetration distance into the virtual object (0.03m). We repeated these steps 40 times (both for the positive condition and for the negative condition) as the number of participants in the experiment.

Fig. 3(d) presents the results of the simulation of the two experimental stretch conditions. Similar to the experimental results, the simulation results show a wide range of perceptual underestimation and overestimation effects for both the negative and the positive conditions. Albeit, in most of the cases, both the negative and the positive artificial stretches increased the perceived stiffness. Moreover, and most importantly, the perceived stiffness due to the positive stretch was consistently higher than the perceived stiffness due to the negative stretch.

### D. Grip Force Control

The grip force results revealed an increase in the applied grip force as a result of the artificial skin-stretch in both skin-stretch conditions. Moreover, the increase in the applied grip force during trials with positive stretch was greater than the increase in the applied grip force during trials with negative stretch. Fig. 4(a-c) show the evolution of the different grip force metrics with repeated interactions for the four different conditions. Surprisingly, already from the first interaction with the virtual object, the grip force of the positive stretch session was higher than the grip force of the negative stretch session in all the grip force metrics, but was identical between the control and the artificial stimulation conditions in each session. Because our interest is in the added effect of the artificial stretch, we focus here on the differences between the grip force values in the positive and negative artificial stretch conditions and each of their control conditions, presented in Fig. 4(d-f), and report the results of the statistical analysis on these differences as dependent variables.

**Fig. 4.**
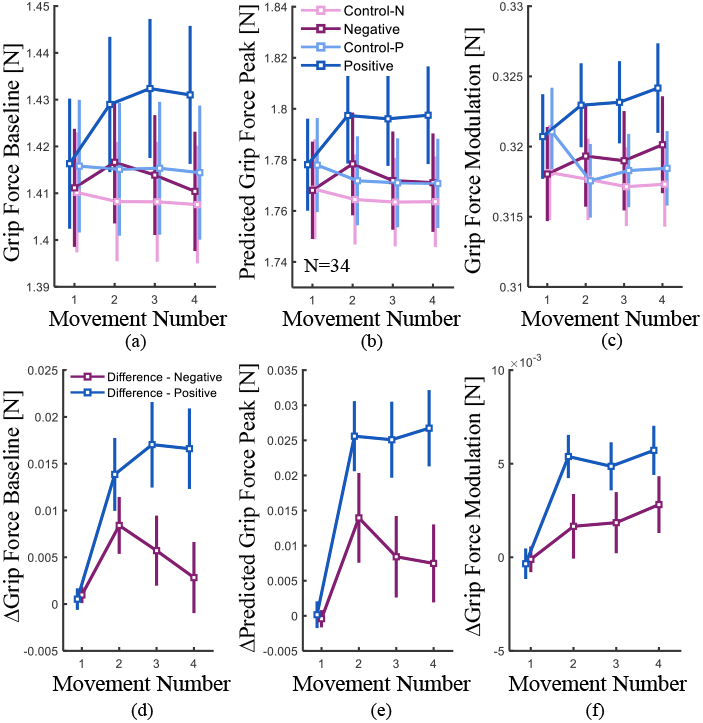
The effect of positive and negative artificial skin-stretch on grip force control. (a) The grip force at contact, (b) the predicted grip force peak, and (c) the grip force modulation, all as a function of the movement number within a single trial. The light purple and light blue traces represent trials without artificial skin-stretch (controls), and the dark purple and dark blue traces represent trials with artificial skin-stretch. The difference between the effect of positive and negative artificial stretches and their controls on (d) the grip force at contact, (e)the predicted grip force peak, and (f) the grip force modulation, all as a function of the movement number. The purple symbols and lines represent the negative stretch session, and the blue symbols and lines represent the positive stretch session. Each symbol is the average value for all the participants (N=34), and the vertical lines represent the standard error.

Fig. 4(d) presents the Δ*grip force baseline* as a function of the movement number. Overall, the increase due to positive stretch was larger than due to negative stretch (rm-General Linear Model, main effect of ‘stretch condition’: *F*_(1,231)_ = 15.05; p = 0.0001). In addition, the Δ*grip force baseline* of both the positive and the negative stretch conditions rapidly increased in the second interaction, but then their patterns of change diverged (rm-General Linear Model, main effect of ‘movement number’: *F*_(3,231)_ = 6.76; p = 0.0002; interaction between ‘stretch condition’ and ‘movement number’ variables: *F*_(3,231)_ = 2.68; p = 0.0474). Specifically, due to the additional positive stretch stimulation, following the immediate increase after the first interaction with the virtual object, the Δ*grip force baseline* of the positive stretch remained relatively constant throughout the remaining probing movements. This was supported by the planned t-tests: there was a significant difference between the first and the second, and the first and the last movements of the positive stretch condition (*t*_P1−2 (231)_ = 3.43, *p*_corrected_ = 0.0014; *t*_P2−4 (231)_ = 0.70, *p*_corrected_ = 0.4791; *t*_P1−4 (231)_ = 4.14, *p*_corrected_ = 0.0001). In contrast, the immediate increase in the Δ*grip force baseline* of the negative stretch condition was followed by a rapid decrease after the second interaction. However, this effect was not supported by significant differences between the first and the second, the second and last, and the first and the last movements in the negative stretch condition (*t*_N1−2 (231)_ = 1.91, *p*_corrected_ = 0.1707; *t*_N2−4 (231)_ = 1.43, *p*_corrected_ = 0.3055; *t*_N1−4 (231)_ = 0.47, *p*_corrected_ = 0.6324).

Fig. 4(e) presents the Δ*predicted grip force peak*, and exhibits similar trends to those of the *grip force baseline*, (rm-General Linear Model: main effect of ‘stretch condition’: *F*_(1,231)_ = 21.40, *p* < 0.0001; main effect of ‘movement number’: *F*_(3,231)_ = 12.24, *p* < 0.0001). However, contrary to the Δ*grip force baseline* analysis, the interaction between ‘stretch condition’ and ‘movement number’ was not statistically significant (*F*_(3,231)_ = 2.53; p = 0.0581). Therefore, the planned t-tests were performed on the two stretch conditions together. The planned t-tests revealed that there was a significant difference between the first and the second movements (*t*_1−2 (231)_ = 5.41, *p*_corrected_ < 0.0001), and between the first and the last movements of the negative and the positive stretch conditions together (*t*_1−4 (231)_ = 4.68, *p*_corrected_ < 0.0001). Here, also, we did not find a difference between the second and the last movements (*t*_2−4 (231)_ = 0.72, *p* = 0.4676). This supports the conclusion following the Δ*grip force baseline* analysis that both stretch conditions increased the overall grip force that participants applied, and that the increase due to the positive stretch was larger. However, the patterns of change between the Δ*grip force baseline* and the Δ*grip force modulation* were different.

Fig. 4(f) presents the evolution of the Δ*grip force modulation* with repeated interaction. As with both of the previous metrics, the increase in the Δ*grip force modulation* of the positive stretch was greater than the increase of the *grip force modulation* of the negative stretch (rm-General Linear Model, main effect of ‘stretch condition’: *F*_(1,231)_ = 8.52, *p* = 0.0039). However, in contrast to the Δ*baseline grip force* and to the Δ*predicted grip force peak*, after the first probing movement, the Δ*grip force modulation* consistently increased with repeated interactions. Fig. 4(f) shows that both the positive and the negative stretch conditions exhibited an increase with repeated interactions with the virtual object (rm-General Linear Model, main effect of ‘movement number’: *F*_(3,231)_ = 6.22, *p* = 0.0004). The planned t-tests revealed that there was a significant difference between the first and the second movements (*t*_1−2 (231)_ = 3.28, *p* = 0.0023), and between the first and the last movements (*t*_1−4 (231)_ = 3.94, *p* = 0.0003) of the negative and the positive stretch conditions together, but not between the second and the last movements (*t*_2−4 (231)_ = 0.65, *p* = 0.5140).

## IV. Discussion

In this study, we examined how the addition of artificial skin-stretch, in the same direction as, and in the opposite direction to, the kinesthetic load force, affect stiffness perception and grip force control during interactions with elastic objects. Our results suggest that, on average across participants, adding artificial skin-stretch in either direction creates the illusion of interacting with a stiffer object. However, this illusion was consistently greater during the interaction with the positive stretch in comparison to the negative stretch. In addition, while the individual effects of most of the participants were qualitatively consistent with these average trends, we observed high variability in the individual effects, including several participants for whom artificial stretch decreased the perceived stiffness. Nevertheless, for most of the participants, the positive artificial stretch led to a higher perceived stiffness than the negative artificial stretch. We suggested a computational model based on the non-uniform distribution of the mechanoreceptors’ preferred direction to explain these results. The model explained all the major aspects of results: the average trends, the high variability in the individual effects, and the heterogeneity in the direction of the individual effects. Positive and negative artificial skin-stretch also led to an increase in the applied grip force. Similar to the perceptual results, positive skin-stretch led to a greater increase in the applied grip force in comparison to the negative skin-stretch.

We reproduced our previous results [27], [29] of augmentation of perceived stiffness due to the positive stretch. This effect is consistent with previous studies, which reported that adding artificial tactile feedback to kinesthetic forces augments the perception of friction [25], stiffness [27], and force [44]. Contrary to our hypothesis, we found that the negative stretch also increased the perceived stiffness for the majority of the participants. However, examination of the individual results revealed two important findings. The first is that there is a large variability across participants in the perceptual effects; our large sample size makes us now confident that this variability also includes participants who exhibit the opposite effect (reducing the perceived stiffness due to artificial stretch). The second is that the increase in perceived stiffness caused by the negative stretch was consistently lower than that caused by the positive stretch, regardless of the effect of the positive stretch. Additionally, the significant difference between the effects of the positive and negative stretch implies that participants probably did not treat the artificial skin-stretch as an indication of a more slippery contact surface, but rather as perceptually-relevant information.

The only previous study that investigated the effect of negative artificial skin-stretch found a large between-participants variability in the perceptual effects, and did not reach a decisive conclusion [30]. Their study included only 12 participants; some of the participants showed a decrease in the perceived stiffness as a result of the negative stretch, and some of them showed an increase. These results led them to conclude that it would not be possible to decrease the perceived stiffness by rendering negative artificial skin-stretch. Our results support this conclusion and even show that it is possible to increase the perceived stiffness by rendering negative artificial skin-stretch. In their study [30], participants also received visual feedback, which may explain the even larger inter-subject variability compared to our results.

The perceptual results of this study shed light on the unique characteristics of the human sensory integration mechanism involved in the perception of stiffness. Contrary to expectations, reversing the direction of the artificial skin-stretch, which created an incongruent direction with the perceived force, did not reverse the perceptual bias, but only reduced the bias in comparison to the bias observed with congruent artificial skin-stretch. These results challenge the traditional sensory integration scheme, which suggests that the perception of force is a result of combining information from multiple sensory modalities [45]. Our previous study [27] that used positive artificial skin-stretch supports this idea by demonstrating that amplified positive artificial skin-stretch led to an enhanced perception of stiffness. Furthermore, adding visual feedback that was congruent with the force feedback, but not with the amplified artificial skin-stretch, reduced this effect [29], providing further evidence for the traditional sensory integration idea. According to the traditional sensory integration idea, if the direction of artificial skin-stretch is reversed, the resulting estimated stiffness would also be reversed compared to the estimated stiffness obtained from a positive skin stretch, or to be ignored due to a lack of reliability. However, the results of the current study show that negative artificial skin-stretch can also generate a perceptual bias similar to the perceptual bias produced by artificial positive skin-stretch. This outcome cannot be explained by a weighted sum of information originating from multiple sensory modalities.

We propose a computational model which may explain the perceptual results of both negative and positive artificial skin-stretch effects. This model is based on two previously reported characteristics of mechanoreceptors and tactile afferents: their non-uniform distribution across the finger pad, and the fact that the tactile afferent neurons have a preferred direction [34], [36]. The results of our simulation showed that in most cases, the two simulated skin-stretch conditions augment the perceived stiffness, and that the perceived stiffness due to the negative stretch was consistently lower than that caused by the positive stretch. These observations are consistent with our experimental results, and therefore the model appears to describe the results well. However, our model failed to accurately predict the sizes of the effects we observed in the experiment. This simulation contained only one degree of freedom: the angle between the direction of the mechanoreceptors’ preferred direction and the direction of the stretch stimulation. We believe that an additional degree of freedom, the amplitude of each of the mechanoreceptors, could have extended the range of effects on the perceived stiffness. However, a model with a larger number of degrees of freedom would require validation with more behavioral conditions. This being said, our simulation predicted the PSE values of the negative and the positive artificial skin-stretch relatively well.

This model can also provide an additional explanation to the between-participants variability observed in the perception effect in previous studies [27], [28], [30] and in this study.

It was previously suggested that this variability could stem from differences between participants’ skin properties [46], [47] or the way that participants held the device [28]. Interestingly, all prior studies (including our own) [27]–[29], [48] interpreted this variability as different levels of sensitivity to stretch, and assumed that some participants are capable of completely ignoring this stimuli. However, this study, which had a larger sample size and stimulation in different directions, allowed us to propose our model and offer an additional explanation for the variability. Participants were instructed to hold the skin-stretch device using their thumb and index finger, while placing the center of the finger pads on the tactors. The outer areas of the finger pads were in contact with the outer shell surrounding the tactors. Additionally, participants were instructed to keep their fingers horizontal with respect to the table. However, we did not constrain the position of the fingers to these locations, and the fingers could move slightly, rather than retaining an exact position throughout the experiment. Therefore, the orientation of the fingers holding the device could vary between trials and between participants. We believe that these different orientations may lead to different magnitudes of stimulation based on the preferred directions of the mechanoreceptors.

Beyond the perceptual effects, we were also interested in exploring the effects of artificial skin-stretch in different directions on the control of grip force. We found that during the first interaction with the object, participants applied a higher *grip force baseline* in the positive skin-stretch session compared to the *grip force baseline* in the negative skin-stretch session. Additionally, already in the second interaction, participants increased the *grip force baseline* in the positive skin-stretch trials, and it remained high throughout the interaction with the object. In contrast, although it appears that the *grip force baseline* of the negative stretch increased in the second interaction and then decreased, these findings did not reach statistical significance. Therefore, it is difficult to conclude whether it is a matter of small effect sizes (albeit our sample size was of a reasonable size) or that the *grip force baseline* of the negative stretch did not change with repeated interaction. These results are not consistent with previous studies [10], [24], [48], which showed a decrease in the *grip force baseline* with repeated interactions with an elastic object. These studies attributed this finding to an increase in the certainty regarding the forces, which resulted in a reduction in the safety margin.

As we observed no decrease in the *grip force baseline* with repeated interactions during the negative stretch session, and an increase during the positive stretch session, we argue that there was a general effect of uncertainty on the control of grip force throughout the entire experiment. This uncertainty may have caused participants to preserve a high safety margin, which was higher during the positive stretch session. The general effect of uncertainty can also explain why the *grip force baseline* of the positive stretch session was higher than the *grip force baseline* of the negative stretch session from the first interaction with the object. This view corroborates the findings of our previous studies that showed that uncertainty caused participants to preserve a high safety margin throughout interactions with objects [27], [49].

The increase in the *grip force modulation* during the positive stretch session was greater than the increase in the *grip force modulation* during the negative stretch session. This increase can be explained by an increase in the perceived load force caused by the skin-stretch stimulation [28], [44]. It is possible that increased perception of load force affected the predictive modulation, and therefore participants increased the *grip force modulation*. Additionally, participants may have interpreted the negative skin-stretch as a low positive skin-stretch gain, and therefore applied lower grip force. This interpretation is consistent with our previous work [27], in which we showed that the *grip force modulation* increased with an increase in the skin-stretch gain. In contrast to [27], we did not find a gradual increase in the *grip force modulation* with repeated interactions. However, we believe that the main reason we did not reach statistical significance is the amount of artificial skin-stretch. Therefore, increasing the skin-stretch gain may cause participants to gradually increase the *grip force modulation* with repeated interactions, as in [27].

It is important to note that the baseline grip forces were higher, and the effect sizes were smaller, compared to our previous study [27]. These two observations are likely related; when the baseline grip force is higher, it becomes less critical to modulate the grip force in anticipation of load forces, as the safety margin is already maintained by the baseline grip force. Similar effects are reported in the absence of force feedback [50], or in grasping of virtual deformable objects [51].

Studying how skin-stretch in different directions affects perception and grip force can contribute both to understanding neural processes and to the development of force displaying technologies. For example, comparing the effects of artificial skin-stretch applied in different directions may shed light on how artificial skin-stretch is interpreted by the nervous system. Additionally, systems that apply artificial skin-stretch to the finger pad have a limited range of movement [28]. Applying skin-stretch in different directions can facilitate the presentation of larger ranges of information in force displaying systems such as teleoperated robotic systems.

Understanding how tactile and kinesthetic information are integrated by healthy individuals when forming stiffness perception and grip force control is important for further elucidating how our brain processes information from the external world via the sense of touch [52]. Additionally, it has many practical implications for improving the design of tactile interfaces in a variety of sensory substitution and augmentation applications such as prosthesis [53], teleoperation [54], and robot-assisted surgery [26]. Our computational model could be an effective design tool for these applications, enabling first the prediction of the perceptual effects in a simulation before proceeding to studies with human participants. However, before this would be possible, future studies that test the reliability of our proposed computational model are needed.

## V. Conclusions

In this work we examined how the addition of artificial skin-stretch, in the same (positive), and in the opposite (negative), directions relative to the kinesthetic load force affect stiffness perception and grip force control during interactions with elastic objects. Our results showed an augmentation in stiffness perception due both to the positive, and to the negative, artificial skin-stretch. However, the perceived stiffness due to the negative stretch was consistently lower than that caused by the positive stretch. Based on these results, we proposed a computational model that predicts the perceptual effect based on the preferred directions of the stimulated mechanoreceptors. Positive and negative artificial skin-stretch also led to a rapid increase in the applied grip force. Similar to the perceptual results, positive skin-stretch led to a greater increase in the grip force in comparison to the negative skin-stretch. This work can be applicable in developing intelligent controllers for robotic hands and wearable finger-grounded tactile haptic devices, and in developing novel technologies for providing haptic information in human-robot physical interactions.

